# Surface Display For Phage Assisted Continuous Evolution: A Platform For Evolving / Screening Nanobodies In Prokaryote Systems

**DOI:** 10.64898/2026.04.03.716437

**Authors:** Fernando F. Mora, Jennifer Brodsky, Ashley Tse, Rory Hoover, Gabrielle Cerna, Benjamin B. Bartelle

## Abstract

Despite >50 years of methods development, specific antibodies are still generated at low throughput and remain in high demand across biotechnology. Most biologics and immunoprobes are monoclonal antibodies, developed using a combination of inoculating animals with a target antigen, engineered candidate libraries, and multiple rounds of selection using phage or yeast display. Here we introduce a synthetic biology scheme to eliminate the need for nearly all of these steps, by combining Surface display on E. coli and Phage display with the microvirus ΦX174, Assisting Continuous Evolution (SurPhACE). Instead of building libraries for screening, SurPhACE runs a closed evolutionary program. A typical experiment can have 10^11^ mutant candidates under active selection, with complete turnover of the mutant population every 30min, or >5×10^12^ unique mutants per day, using less than 100mL of bacterial culture media. We demonstrate SurPhACE for optimizing a nanobody to a related epitope, and develop novel nanobodies for an arbitrary target using a minimal starting library to establish a proof of concept and identify best practices for this scalable method for generating protein binders.

## Introduction

Binding proteins are small, protein-based entities that can recognize and bind to a particular target in a selective manner^1^. They are a broadly useful tools in research, industry and have an important role in healthcare where “biologics” comprise a significant portion of the new drugs approved by the FDA^2–4^. These can be classified into immunoglobulin-derived: antibodies single-chain variable fragments (scFvs), V_NAR_s, nanobodies and non-immunoglobulin-derived protein scaffolds like designed ankyrin repeat proteins (DARPins), monobodies, affibodies, anticalins and cystine-knot peptide (knottin)^1,5–8^. Generating a functional new binding protein requires a mixed strategy of rational design choices, and Directed Evolution (DE) with a maximal library size. There are a variety of strategies to maximize the mutational space explored while biasing the search to areas with the highest probability of generating a gain of function mutant ^3,7,9–13^.

Immunoglogulin-derived proteins remain an overwhelmingly popular chassis for engineering, largely because they can be derived from the natural hyper-evolution of the vertebrate adaptive immune system^5,14–16^. Immunization of a live animal produces evolved optimized antibody binders that can be coarsely purified as antisera, plasma B cells can be isolated hybridized and screened for optimal monoclonal antibodies, and successful clones can be sequenced for production in other cell lines^17,18^. These methods have been scaled and optimized over decades, but still require an animal facility and months to generate effective binders to a new target^3^.

Phage and yeast display technologies build on animal derived antibodies and our knowledge of their structure function^19,20^. Whether a species derived antibody has a single domain or heterodimerized heavy and light chains, the complementary determining regions (CDR1-3) are responsible for nearly all binding affinity, with CDR3 being dominant and CDR1-2 largely defining allowable motifs^4,21^. Commercial antibody libraries each apply their own strategy to optimize this starting diversity for libraries of >10^9^ ^10,22–24^. Deriving functional antibodies requires an experimental selection strategy, often biopanning, with iterative re-amplification and tuning with saturation mutagenesis of CDR1-3, exploring a total mutational space of >10^10^ variants^23,25,26^. In vitro DE can be performed at the bench by a single operator, remains labor intensive^27,28^.

In contrast to in vitro saturation mutagenesis, Phage Assisted Continuous Evolution (PACE) can potentially generate very large unbiased libraries of unique mutants, which are constantly under selection pressure^29–31^. Here a gene of interest (GOI) is integrated into a phage genome, which subsequently infects host bacteria engineered for a molecularly defined selection condition and accelerated mutation rate. Because mutations are random, they are sparsely distributed across both the GOI, the phage genome that carries it, and the host bacteria, which must be continually replenished with fresh stock to retain viability^32,33^. The unbiased approach of PACE is advantageous, even necessary for proteins that lack extensive characterization, but might be a neutral trade-off for library size when evolving defined motifs.

While molecular techniques have proven to be reliable, the raise of artificial intelligence (AI) tools have opened possibilities beyond what evolution can do with protein binders^34–37^. Several novel protein binders have been developed with high levels of affinity by these or similar tools^38–43^. However, limitations like training biases, protein structure prediction and the modeling the CDR3 of nanobodies remain to be a challenge, even for the most advances AI approaches^35,44^. For these reasons molecular approaches are still continuously developed and will stay relevant for the foreseeable future and complement *in silico* based techniques.

Here we develop a method to couple the massive mutational space explored by PACE with rational library design for designing novel binding proteins. Surface Display for Phage Assisted Continuous Evolution (SurPhACE) utilizes a novel engineered coliphage ΦX174, dual phage/bacteria display, and simultaneous positive/negative selection of candidate protein binders. SurPhACE is compatible with selection from in vitro derived libraries and spontaneous library generation with mutational plasmids developed for PACE.

We demonstrate evolutionary principles at work and identify crucial best practice concepts for SurPhACE and DE methods in general. The examples provided utilize the camelid single chain variable domain, or nanobodies, but the method is binding protein agnostic. The goal of the work described is an accessible modular DE platform for benchtop scale binding protein engineering, using only standard molecular biology equipment and constructs available on AddGene.

## MATERIALS AND METHODS

### Media

All experiments were conducted using lysogeny broth (LB) supplemented with the appropriate antibiotics. The lactose induction medium (LIM) had ampicillin (100 ug/ul), chloramphenicol (20 ug/ml) and lactose 10 mM. The arabinose induction medium was made in two versions: one containing ampicillin (100 ug/ul), chloramphenicol (20 ug/ml), glucose (0.05%) and arabinose 25 mM, and was used to grow the helper bacteria alone. We will refer to this medium as arabinose induction medium 1 (AIM1). The second version did not have glucose and it was used during the electroporation step to activate MP6 immediately. This will be referred as arabinose induction medium 2 (AIM2). Glucose-only supplemented medium with antibiotics (ampicillin with / or chloramphenicol) was used to make glycerol stocks of the helper bacteria and LB – agar plates (1.5% agar)

### Plasmids design and cloning

All the plasmids used in this work were derived from a modified pRSET backbone. To construct the helper plasmids, 1 Gblock DNA fragment of the target protein with an in-frame upstream BAN anchoring motive attached were ordered from Integrated DNA Technologies (IDT). The *Bacillus Anthracis* N terminal peptide (BAN) came from the exosporal protein of *Bacillus anthracis* and its purpose was to display the target protein on the outer membrane of *E. coli*^45^. The H gene was isolated from New England Biolabs (NEB) ΦX174 RF I DNA. PCR was used to amplify and add the linkers to the ends of the DNA parts for the cloning reactions. All PCR and cloning reactions were performed using NEB High Fidelity Q5 Master Mix 2X and NEBuilder HiFi DNA Assembly Master Mix respectively according to the manufacturer instructions (Figure 1, Supplementary table 1. All genes were placed under the control of a modified *tac* promoter which was turned into a constitutive promoter by removing the Lac operator downstream of the sequence to ensure availability of the H protein to the bacteriophages. After the cloning procedure, 1 ul of the assembly reaction was used to transform NEB® Stable Competent *Escherichia. coli*, then 50 μl of a 1/10 diluted mixture was spread onto agar plates containing ampicillin and incubated at 30 °C overnight. The next day, 6 colony candidates were used to prepare 3 ml LB cultures supplemented with ampicillin which were PCR screened for the presence of the insert of the right size. The positive cultures had their plasmid extracted using NEB Monarch Spin Plasmid Miniprep Kit following the manufacturer instructions. Subsequently, the plasmids were sent for sequencing by Plasmidsaurus using Oxford Nanopore Technology with custom analysis and annotation. The decoy plasmid was made in a similar manner, but the *tac* promoter was left intact and the H gene was omitted from the plasmid.

**Figure 1.**
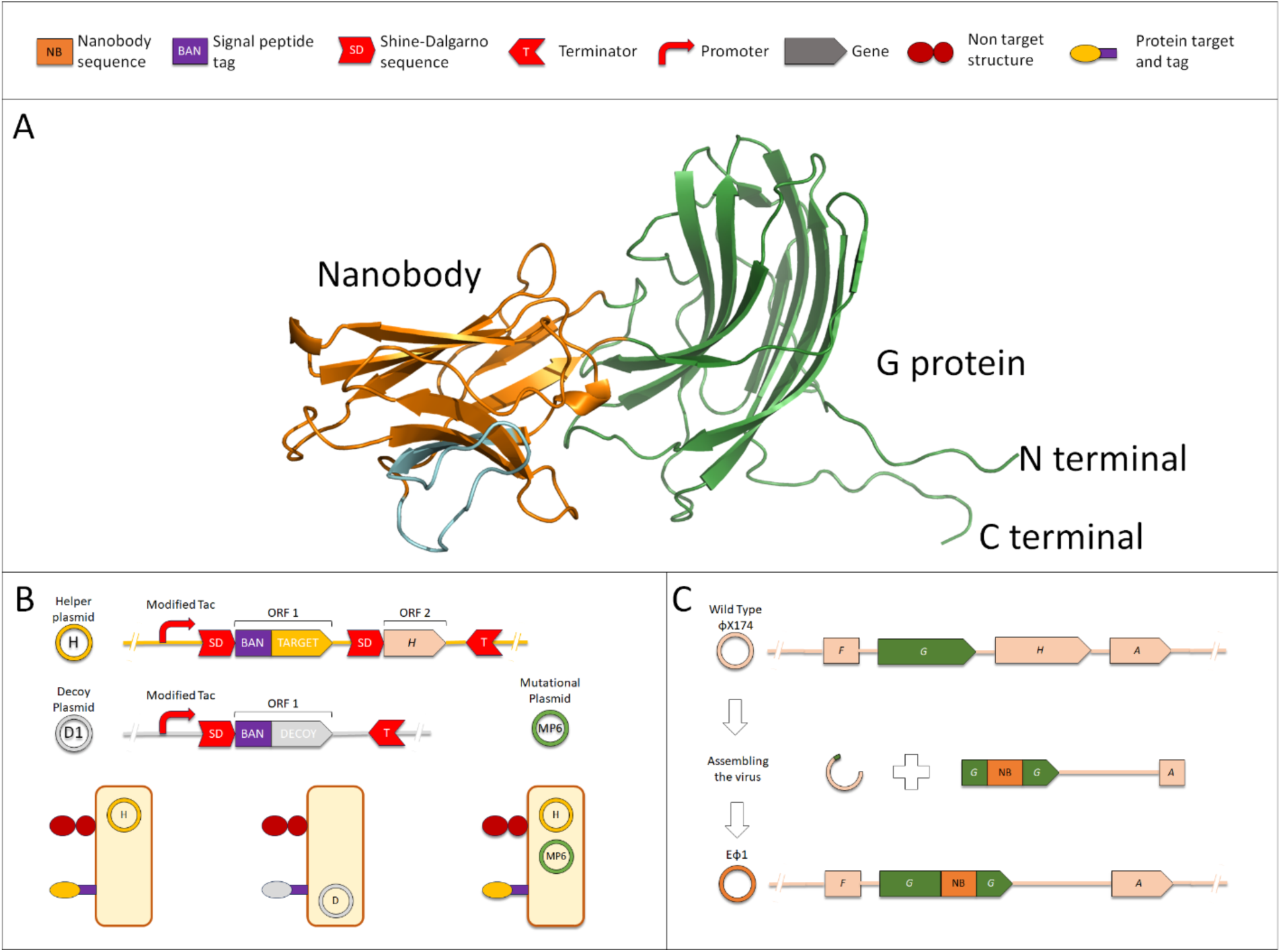
Depiction of all the components of SurPhACE. **A.** AlphaFold model of the complete fusion spike protein, colors match the cartoon drawings. **C.** Graphical description of the cloning process of the engineered phage EΦ1, the wild type genome has its H gene removed.

Three proteins targets were chosen and inserted into different helper plasmids backbones. The first target was the 28-288 aa region of the Fc gamma receptor IIa (gene name: FCGR2A) and it was used to test the components and standardize the technique conditions. The other two targets, the globular protein progestagen associated endometrial protein (gene name: PAEP) and the 1-142 aa soluble portion of the interleukin 20 receptor subunit alpha (gene name: IL20RA), were used to test SurPhACE capabilities to evolve and screen for nanobody candidates.

The mutational plasmid used was MP6^46^ and it was purchased from Addgene (catalog number: #69669) and the *E. coli* harboring it were kept in medium supplemented with glucose at 25 mM according to the authors suggestions. These bacteria had their plasmids extracted using NEB Monarch® Spin Plasmid Miniprep Kit according to the manufacturer instructions. The complete helper bacteria contained some variant of the helper plasmid with the MP6.

### ΦX174 engineering

The phage used to carry the nanobody was ΦX174, a single stranded DNA virus of the family Microviridae. The phage DNA template used was NEB ΦX174 RF I DNA. The nanobody (M2e-VHH-23m / nanobody-αFcR2B) was obtained from De Vlieger *et al* 2019^47^ and the DNA sequence was inserted after cysteine 312 in the G gene (Figure 1A). For the experiments depicted in Figure 2, we opted to use an innovative version of PCR assembly to build the engineered ΦX174 that we will be referring as EΦ1. A Gblock containing part of the G gene (48 bp), the nanobody and filler DNA with part of the A gene (63 bp) was ordered from IDT. The filler DNA was a random sequence (luciferase) that was put in there to fulfill the phage genome size requirements^38^.

**Figure 2.**
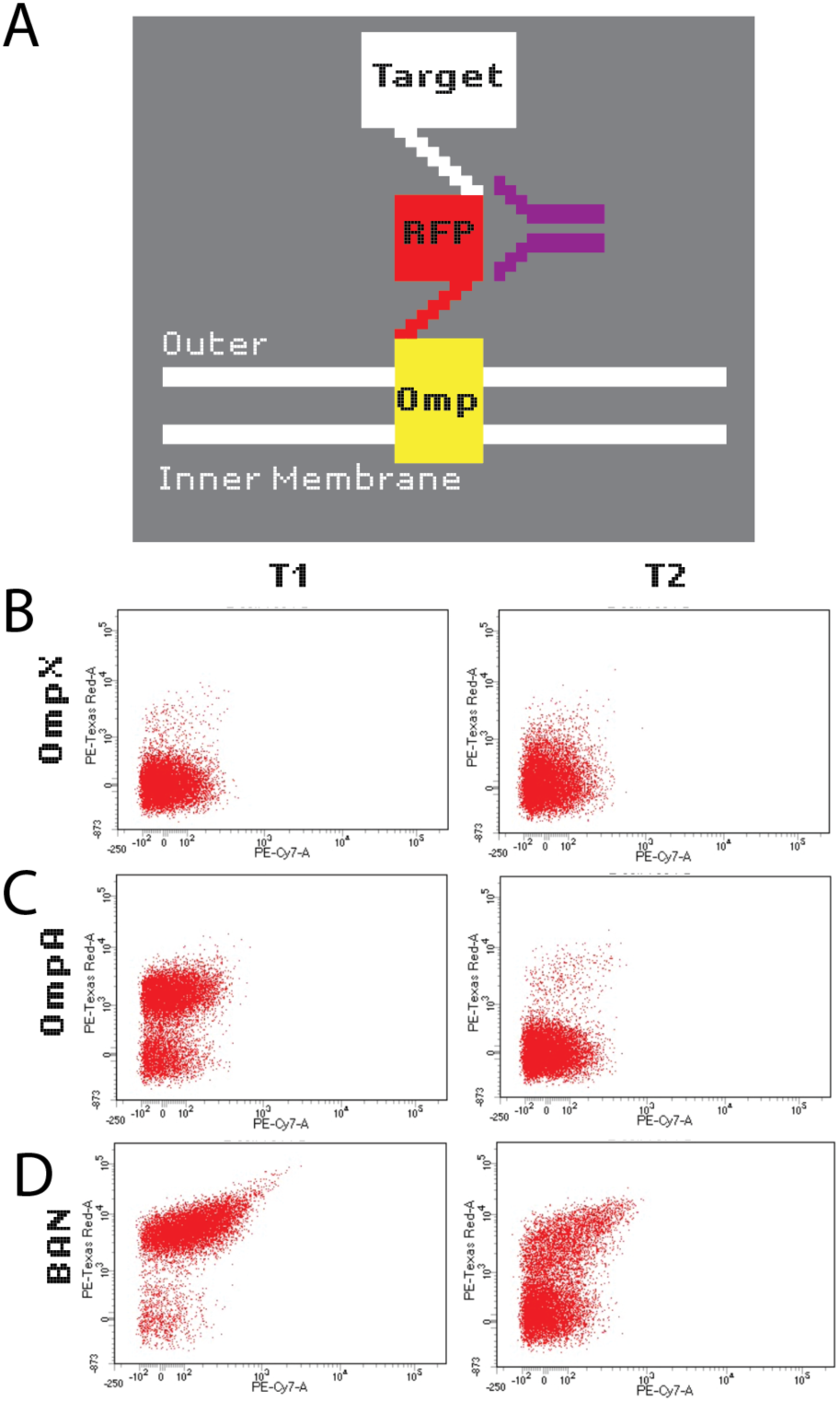
Flow cytometry experiments using 3 different tag systems. **A.** Schematics showing the general tagging procedure, depicting Omp as an example. The fusion protein: tag, Fluorescence Protein and the target is displayed on the outer membrane of E. coli. **B-D.** Identical setup but different tag proteins, for two different targets, T1 and T2. **B.** OmpX. **C.** OmpA. **D.** BAN

The procedure to build EΦ1 is described as follows: the ΦX174 genome was linearized by PCR amplification using the primers P-Inv-5 and P-Inv-6 for 45 cycles. The product was gel purified using NEB Monarch Spin DNA Gel Extraction Kit and the DNA concentration of the eluate was measured using a Nanodrop. The linearized phage solution was mixed in a 1:10 mass ratio (50 ng of linear ΦX174) with the Gblock, the later worked as a megaprimer in a PCR reaction for 45 cycles. All PCR reactions were done using NEB High Fidelity Q5 Master Mix 2X reagent according to the manufacturer instructions unless stated otherwise (98°C, 30 s | (98°C, 5 s \_72°C, 2 min) X 45 | 72”C 10 min). The PCR reaction was then precipitated with 1/10 volumes of sodium acetate 3M pH 5.2 and 2.5 volumes of ethanol 100% and the solution was mixed by inverting the tube 6 to 10 times and placed in the freezer at -20°C overnight. The next day the solution was centrifuged, the supernatant was discarded and 200 ul of 70% ethanol was added without disturbing the pellet and let it incubate at room temperature for 5 min. Immediately after, the ethanol was removed and the pellet was resuspended in 50 - 100 ul of deionized sterile water. The phage prepared this way could be stored indefinitely in its precipitated form or resuspended in water as long as the freezing / thaw cycles do not exceed 10 times.

### Artificial CDR3 library

A YNN type library was made by randomizing the complementary determining region 3 (CDR3) of the nanobody. To do this, a 100 nt long DNA oligo was ordered from IDT. This oligo had 20 nt overlaps at both ends and 60 nt long randomized region. In order to incorporate this modified CDR3 into the ΦX174 genome, two additional parts were needed. One part includes the rest of the nanobody, located upstream of the CDR3 sequence, with an overlapping region (30 bp) that matches the genomic G gene at its 5’ end. This part was amplified from the Gblock used as megaprimer for the construction of EΦ1 using primers P2.3 and Part-2-Con-R. The second DNA part correspond to the downstream region containing the remaining of the 5’ nanobody sequence, G gene and the DNA filler region. The 3’ end of this sequence had an overlapping region (63 bp) with the 5’ end of the A gene of ΦX174. The primers used to make this part were P2.4 and Part-2-3, the template for the PCR reaction was also EΦ1 megaprimer. Finally, the ΦX174 genome was linearized using the primers P-Inv-5 and P-Inv-6. All PCR reactions were done using NEB High Fidelity Q5 Master Mix 2X reagent according to the manufacturer instructions (98°C, 30 s | (98°C, 5 s \_ 65°C, 30 s _/ 72°C, varies) X 35 | 72”C 10 min). The new engineered phage library was created by putting these 4 parts together using a NEBuilder HiFi DNA Assembly procedure according to the manufacturer instructions for single stranded oligos with some minimal modifications described as follows: a 50 mM NaCl solution was used instead of water to further stabilize the oligos and the molar ratio between the linearized genome and the other two parts surrounding the oligo was kept the same as the standard protocol (Figure 5A). The reaction was precipitated and resuspended in deionized sterile water as previously described. These phage libraries will be referred as EΦ2.

### *E. coli* bacteria preparation

*Escherichia coli* (Migula) Castellani & Chalmers, C-strain, was purchased from ATCC® (catalog number: 13706) and was used to prepare electrocompetent cells according to the protocol in this kit: Ph.D™.-12 Phage Display Peptide Library Kit, modified for 50 ml cultures. The sequence-verified helper plasmids and MP6 plasmid were co-electroporated using a PrecisionPulse^TM^ ECM 630 electroporator (conditions: 2500 V, 200 Ω, 25 μF) into electrocompetent *Escherichia coli* C-strain to make the complete helper bacteria (Figure 1C). For the decoy bacteria, an identical procedure was followed, however a modified helper plasmid without a target nor a H gene was used instead (Figure 1B). These bacteria were used to prepare electrocompetent bacteria for later use and storage.

### Semicontinuous evolution experiments

The phage genomes (EΦ1 / EΦ2) were electroporated into electrocompetent bacteria depending on the experimental needs (Figure 2B, Figure 5B). After electroporation, 1 ml of induction medium was added and the contents were mixed by pipetting. After mixing, the mixture was pipetted out of the cuvette and transferred immediately into regular test tubes (16 ⌀ mm x 100 mm) containing induction medium. Subsequently, the cultures were properly labeled and placed upright in a shaking incubator (30°C at 250 rpm) for the remainder of the experiment. For more details refer to Figure 2 and Figure 5 for each experiment particular conditions

The following days, a fixed volume of medium and old bacteria was removed from the cultures and replaced by the same volume of helper bacteria and fresh medium. The fresh bacteria were prepared by streaking glycerol stocks of the helper bacteria on LB plates (1% agar, ampicillin (100 ug/ul), chloramphenicol (20 ug/ml) and glucose (25 mM)). The next day a colony was used to seed a culture with LB medium (ampicillin (100 ug/ul), chloramphenicol (20 ug/ml), glucose (0.05%) and arabinose 25 mM), the culture was placed in the shaking incubator (30°C at 250 rpm) and let it grown for 3h or until visible growth was seen (OD ≈ 0.1 - 0.3) to ensure MP6 did not activate prematurely and prevent helper bacteria from dying due to the toxic nature of the helper and mutational plasmids. For the artificial library experiments, the decoy bacteria were prepared from an overnight culture grown in LIM medium. One ml was used at the following time points: 0h and for condition HP (high pressure) at 72h, where 1 ml of the culture was centrifuged (8000 RPM), the old medium discarded and resuspended in AIM2. The high concentration of decoy present at 0h and 72h (the later for high pressure conditions) were done to remove as much unspecific binders as possible. For the other time points 100 ul of decoy bacteria resuspended in AIM2 was used instead and for condition LP at 72h (low pressure). The occurrence of the phages in the cultures was checked by PCR (GoTaq® Green Master Mix 2X, according to manufacturer instruction) using 1 ul of the culture and the primers P1.1 and P1.3 and they were designed to land on the G gene contiguous to the nanobody, the positive control (Wild Type phage DNA) and negative control were controls for the PCR only. For more details about any of the particular experiments refer to Figure s 2 and 4.

### Flow Cytometry assays

*E. coli* C strain was transformed with a decoy plasmid containing a FLAG tag attached to FcγRIIA in frame with a BAN signal. The bacteria were grown in LIM medium and collected moments after reaching mid-log phase. The general procedure is described by Dietrich^48^ and the flow cytometer used was a Attune NxT Flow Cytometer

### Sequencing and data analysis

We used Next Generation Sequencing (NGS) service provided by the Biodesign Institute at ASU to sequence the phage DNA containing the nanobody CDR3 which correspond to the experiment in Figure 2C. The library was made according to Buenrostro *et al* 2015 and the region amplified was 250 bp long. The paired-end fastaq files were analyzed using RStudio, the package Rfastp was used for the quality control and reads merging steps. The package blaster was used to search for mapping out the mutations and manipulating the CDR3 sequences. Other packages like, Biostrings, dplyr, DECIPHER and bioseq were used to further process the DNA data and the CDR3 variant libraries.

In the case of the synthetic CDR3 library, we made use of Plasmidsaurus’ custom project services using Oxford Nanopore Technology to sequence the variant populations at 3 different time points for a total of 4 samples per experiment (Figure 5B). The DNA library was made in the same way as with the NGS sequences and the sequence amplified was the complete nanobody with a small fragment of the G gene at the 5’ end that was used as a frame of reference for the translation of the sequences. The data generated was analyzed using RStudio in a similar way, but Rfastp was only used to check the quality the fastaq files. The code is provided in the supplementary material. The CDR3 variant library was used to look for the complete nanobody sequence attached to each CDR3 variant to analyze the mutations outside the CDR3. A fragment of the G gene in the 5’ upstream region of the nanobody was used as a point of reference for the translation of the variant sequences to differentiate truncated products from complete nanobody proteins. The CDR3 amino acid sequences were analyzed separately using hierarchical cluster analysis in R using a distant matrix calculated using the packages Biostrings (StringDist, with BLOSUM62 as the substitution matrix) and the built-in hclust R function and the amino acid patterns found were analyzed assuming a random uniform distribution.

### Protein modeling

AlphaFold server (AlphaFold 3) was used to predict the structure of the different proteins and also the complex of the nanobody candidates and their targets. ipTM, pTM and to a lesser extent plDDT values were used to determine how well the nanobody candidates might bind and to compare them to actual binding data.

### Mathematical modeling

We designed a mathematical model to describe the population dynamics of the variants and the bacteria present in the cultures. The model has a total of 6 ordinary differential equations (ODE), which includes 1 variable describing the substrate, 2 describing the populations of helper and decoy bacteria and 3 variables describing the phages that bind to different targets. The equations were solved using RStudio and the package deSolve. To simulate evolution, the constants describing binding affinity were altered each time using R code. A diagram showing the interaction of the variables and constants can be seen in Figure 4. The code and a table describing the variables are provided in the supplementary material

## RESULTS

### SurPhACE leverages the biology of Sinsheimervirus ΦX174

Phage display and PACE employ Coliphage M13 to carry genes of interest (GOI) for engineering. M13 infects *E coli* by binding the bacterial flagella, rather than the cell surface, presenting a possible issue for a dual display method. To avoid this, we chose a phage that interacts directly with the cell surface, microviridae ΦX174. In the phage lifecycle, protein G serves as a spike protein, making the first contact with the LPS on the surface of the bacteria, followed by its dissociation and leaving the F protein to make contact with the surface, then the injection of DNA by protein H proceeds^49^. Previous studies identified several sites in protein G as amenable for peptide display^50^. In an AlphaFold model site C312 allowed for correct folding of protein G with the insertion of an entire single chain camelid nanobody and 2 linker regions (**Figure 1A**). The model also suggested the insertion chimera could occlude the binding domain of protein G, suppressing its innate function (**Supplementary Figure 3**).

ΦX174 was the first sequenced, synthetically reconstructed, and refactored genome, demonstrating how amenable it is to engineering. One described caveat is that the genome size is optimized for the phage particle and less tolerant to large insertions or deletions than M13. We fused a nanobody into protein G and excised the entire gene for protein H from the ΦX174 genome, but retained the genome size with random sequences to create what we hypothesized to be a replication defective virus (**Figure 1C**).

### SurPhACE plasmids display proteins on the bacterial cell surface

SurPhACE selection requires direct surface binding of phage displayed antibodies to targets on the cell surface of a bacteria for positive selection, with negative selection against off target binding. This required robust bacterial surface display of arbitrary size protein. We first compared bacterial surface display tags. OmpA and lppOmpX are surface display tags derived from *E. coli* outer membrane proteins, while the BAN tag is derived from a sporulation protein (**Figure 2A**). Using 2 arbitrary proteins (T1 and T2) on 3 different display tags we found that OmpA (**Figure 2B**) and OmpX (**Figure 2C**) showed poor expression and minimal surface display using flow cytometry labeling. Conversely BAN tagged proteins showed consistent labeling correlated to expression level (**Figure 2D**).

### Design and testing of SurPhACE helper and decoy plasmids

We designed a selection scheme for our ΦX174 engineered to be replication defective for lack of protein H and dependent on a displayed nanobody for cell surface binding (**Figure 1C**). For positive selection, we made a multicistronic plasmid containing the BAN displayed target, with a supplemental protein H (**Figure 3A**). To counter nonspecific and off target binding we developed a negative selection plasmid, expressing only the BAN tag, but no target or supplemental genes (**Figure 3B**). Finally, to accelerate evolution, we employed a previously described mutational plasmid.

**Figure 3.**
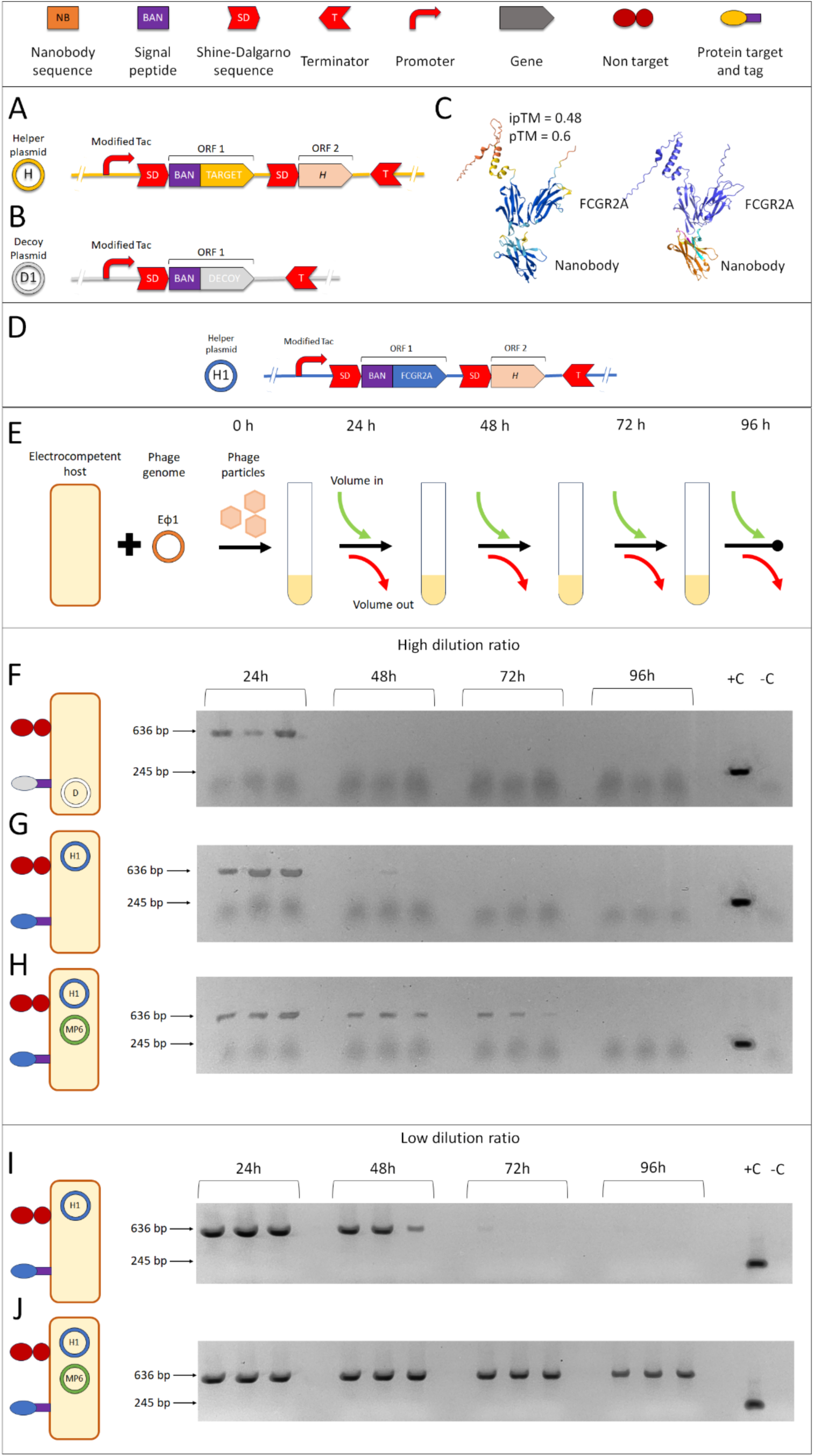
Initialization of SurPhACE components. **A.** Basic layout of the genes in the helper plasmid, this plasmid has the BAN + target and the phage H gene driven by a constitutive promoter and both are transcribed as a polycistronic mRNA. **B.** The decoy plasmid only contains the BAN in frame with a non-target protein, it the genes are transcribed in a identical way as the helper plasmid. **C.** Alphafold hypothetical interaction of the nanobody and the FCGR2A receptor, the 3D model is shown in the Alphafold standard colors showing that the majority of the interacting regions are in blue (plDDT > 90) and cyan (90 > plDDT > 70). **D**. Layout of the helper plasmid 1 (H1) containing the FCGR2A target. **E.** Diagram showing the general setup for the standardization experiments, 2 ul of EΦ1 was electroporated into 100 ul of electrocompetent host bacteria and the contents were immediately poured into a test tube containing AIM1, the phage particles are released into the culture after the host bacteria were lysed. **F.** Gel electrophoresis showing the presence of the EΦ1 phages in the cultures with decoy bacteria at different time points (636 bp band). **G.** EΦ1 phages in the cultures with H1 host bacteria. **H.** EΦ1 phages in the cultures with helper bacteria H1+MP6. The amount of volume of fresh bacteria added and old bacteria removed was 1.5 ml, the total volume of the 3 replicates were kept constant at 3 ml for the 3 experiments (F, G, H). **I.** The experiment depicted in the panel G was modified to be less stringent by increasing the culture volume up to 5 ml and the amount of medium added / removed every day was 1 ml. **J.** Identical setup to (I), but the host bacteria used was the same as (H)

To determine the initial selection conditions for SurPhACE we sought out a candidate nanobody/target pairing with sub-optimal affinity. The surface protein FcγRIIB has a published nanobody and an 90% sequence similarity with FcγRIIA. Computational docking using Alphafold 3 predicted potential affinity between nanobody-αFcR2B and unglycosylated FcγRIIA (ipTM = 0.48), informing a hypothesis that SurPhACE would result in a mutant with increased affinity (**Figure 3C**). To test this, we cloned the extracellular domain of FcγRIIA into our positive selection display/helper plasmid (**H1, Figure 3D**).

To initialize our system, we synthesized a nanobody-αFcR2B-ΦX174 (EΦ1) phagemid and electroporated into E coli expressing our test conditions. Protein H is only required for phage particle formation; thus we expected initial phage replication across all conditions regardless of phage viability. Cultures were refreshed semi-continuously with a set dilution rate of 0.002 h^-1^, low compared to PACE in anticipation of low phage viability (**Figure 3E**).

To test whether our EΦ1 had any starting affinity towards E coli and was capable of replication without protein H, we first tested out SurPhACE using only decoy plasmid D *E coli*. EΦ1 DNA was detectable by PCR at 24h post electroporation of synthetic phagemid, however by 48h no phage genomes remained (**Figure 3F**).

We then tested if our EΦ1 had any starting affinity towards *E coli* and was capable of replication with supplemental protein H. Electroporating phage into H1 *E coli* again resulted in detectable phage genomes by 24h, with signal persisting in one sample to 48h. Phage with H1 did not result in propagation beyond what could be explained by replication in electroporated cells (**Figure 3G**).

In a full test of our system, we electroporated EΦ1 phagemids into E coli with both H1 and MP6. Under these conditions, phage genomes were detectable out to 72h, suggesting propagation was occurring post electroporation, but selection conditions were too stringent for phage survival (**Figure 3H**).

Given our experimental conditions, the only variable that could be changed without choosing a new nanobody/target pair was the washout rate of fresh bacteria. increasing the volume of the culture and reduce the amount of fresh bacteria / medium added, effectively reducing the dilution rate from 0.002 h^-1^ to 0.008 h^-1^. Repeating the electroporation of EΦ1 phagemids into E coli with H1 yielded detectable phage out to 72h (**Figure 3I**) with addition of MP6 extending signal to 96h (**Figure 3J**).

### Phage propagation was not due to conserved mutations in the CDR3 region of EΦ1

The extended survival of phage in H1/MP6 cultures suggested phage propagation, but we could not determine if evolution occurred. To look for mutations, we generated 6 independent replicate libraries of post 168h selected EΦ1 and used next generation sequencing to examine the CDR3 domain that could have conferred survival advantage. CDR3 mutants were rare, comprising less than 1% of the population (**Supplementary Figure 1, Figure 4A**). The number of individuals per variant was also low, with the most abundant mutation only detected in 53 of 36968 viable sequences for replicate 6 (**Figure 4B**). No deletions or insertions were detected in the CDR3 region and the number of substitutions was low, with the most common mutations being synonyms for the current amino acids. These results showed that the role of MP6 in the survival of the phage in these experiments does not require changes in the CDR3 region of the nanobody, but we could not rule out mutations across the rest of the phage genome with this method.

**Figure 4.**
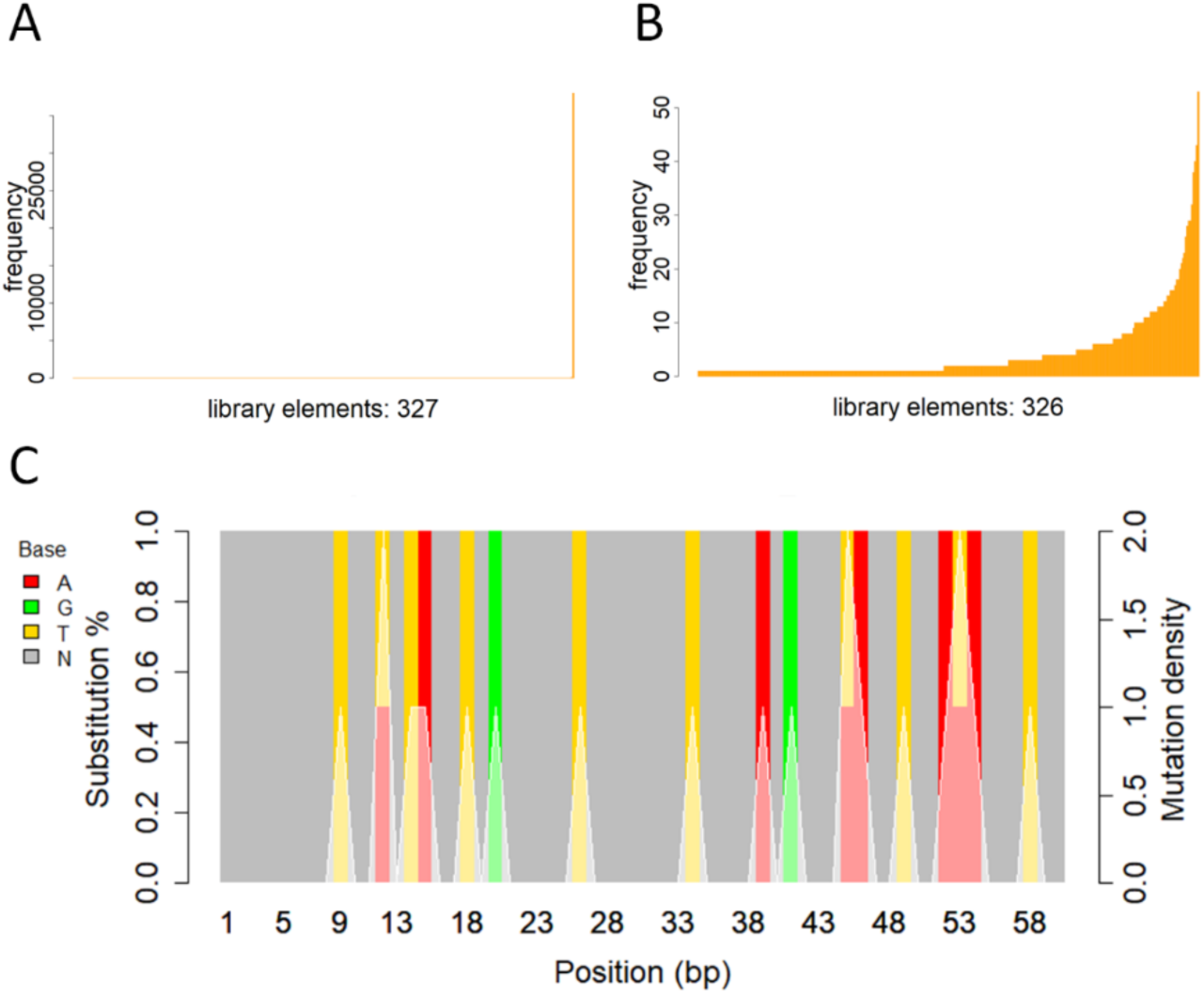
NGS analysis of EΦ1 CDR3. **A**. Number and frequency of CDR3 variants including the variant corresponding to unmodified CDR3. **B**. This graph shows the same information as A, but the variant with the unmodified CDR3 was removed. **C**. Bar plot showing the substitution percentage (colored bars, N stands for unaltered nucleotide) and mutation density (white line) for the 20 most abundant mutated variants.

### Dynamic modeling of SurPhACE based on initialization conditions

For insight into our initial experiments, we generated an ordinary differential equation (ODE) model of our system (**Figure 5**). We initialized our model for a condition where phage survival required mutation and optimal flow rate as we had seen experimentally (**Figure 3**). In the model, an 8.9% gain of function to either infectivity or viability was enough to select out phage at a threshold value (**Figure 5B**). Assuming a 30 min viral life cycle, the complete replacement of WT with a gain of function mutant took less than a day. Above threshold selection levels however, rare mutants could not gain dominance over the population in the timeframe of previous experiment (**Figure 5B**). This aligned with the ratio of variants and non-mutated phages observed in our results (0.008663 ± 0,000441, n = 6, **Supplementary Figure 1**) where the ratio of non-mutated phages and mutated variants were 0.003205 for neutral selection simulations and 0.100310 for positive selection, suggesting the need for a normalization step to enrich rare mutants, or not begin with a single genotype.

**Figure 5.**
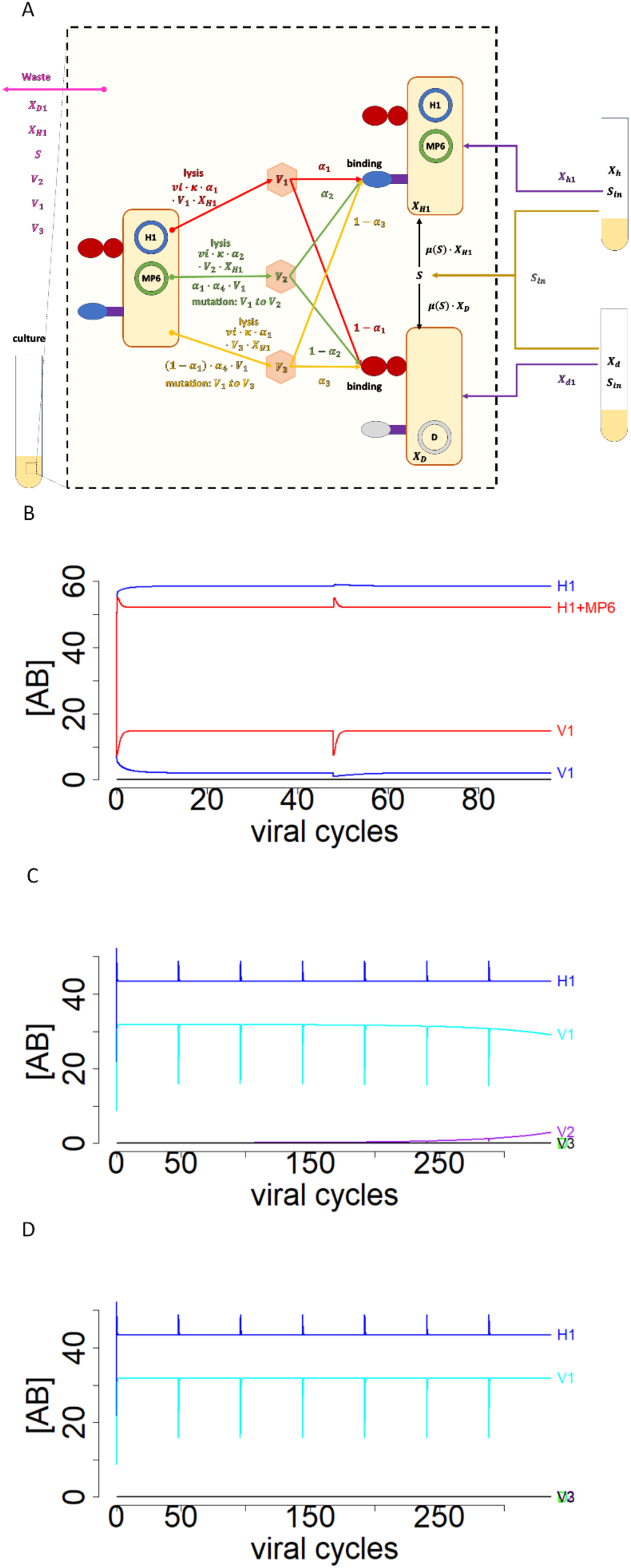
Dynamic model of SurPhACE: A. Scheme showing the variables and possible interactions between them in the culture. The arrows and labels represent the rate of conversion between the variables or the frequency of binding of the phages. B The model was run with arbitrary constants for 2 different phage viability (*vi*) conditions, mimicking the experiments in figure 2D. In the simulation with H1 helper bacteria, a viability of 0.00445 was implemented and for H1 + MP6 the viability was 0.005. C. The model was run again to replicate the results observed in figure 3. The viability was kept at 0.005 and the mutations were set for positive selection. D. The experiment was repeated again but the mutations were set to neutral selection. Simulation Parameters: a1 = 0.96, a2 = a1, a3 = a1, a4 = a4, Dth = 0.5, Y1 = 2, Y2 = 2, Sin = 10, k0 = 2, k1 = 6, k2 = 6, u1 = 9, u2 = 10. Initial conditions: S = 10, H1 = 10, D = 0, V1 = 10, V2 = 0, V3 = 0

### Binding to new targets and CDR3 artificial library

Based on our model, we hypothesized that the propagation of gain-of-function mutations in EΦ1 phage was suppresed by the high abundance of moderate binding variants compared to mutated variants and the inherent basal affinity of the nanobody for FCGR2A. To overcome this, we synthesized a CDR3 mutant library inserted into EΦ1 (EΦ2) (**Figure 6A**). With an expected library size of 10^6^, low binding affinity variants would be rare and compete with each other and worse binders while subject to random mutation. To exclude the initial affinity from CDR1-2, we generated new target helper plasmids to display IL20RA (H2) and PAEP (H3) (**Figure 6B**).

**Figure 6.**
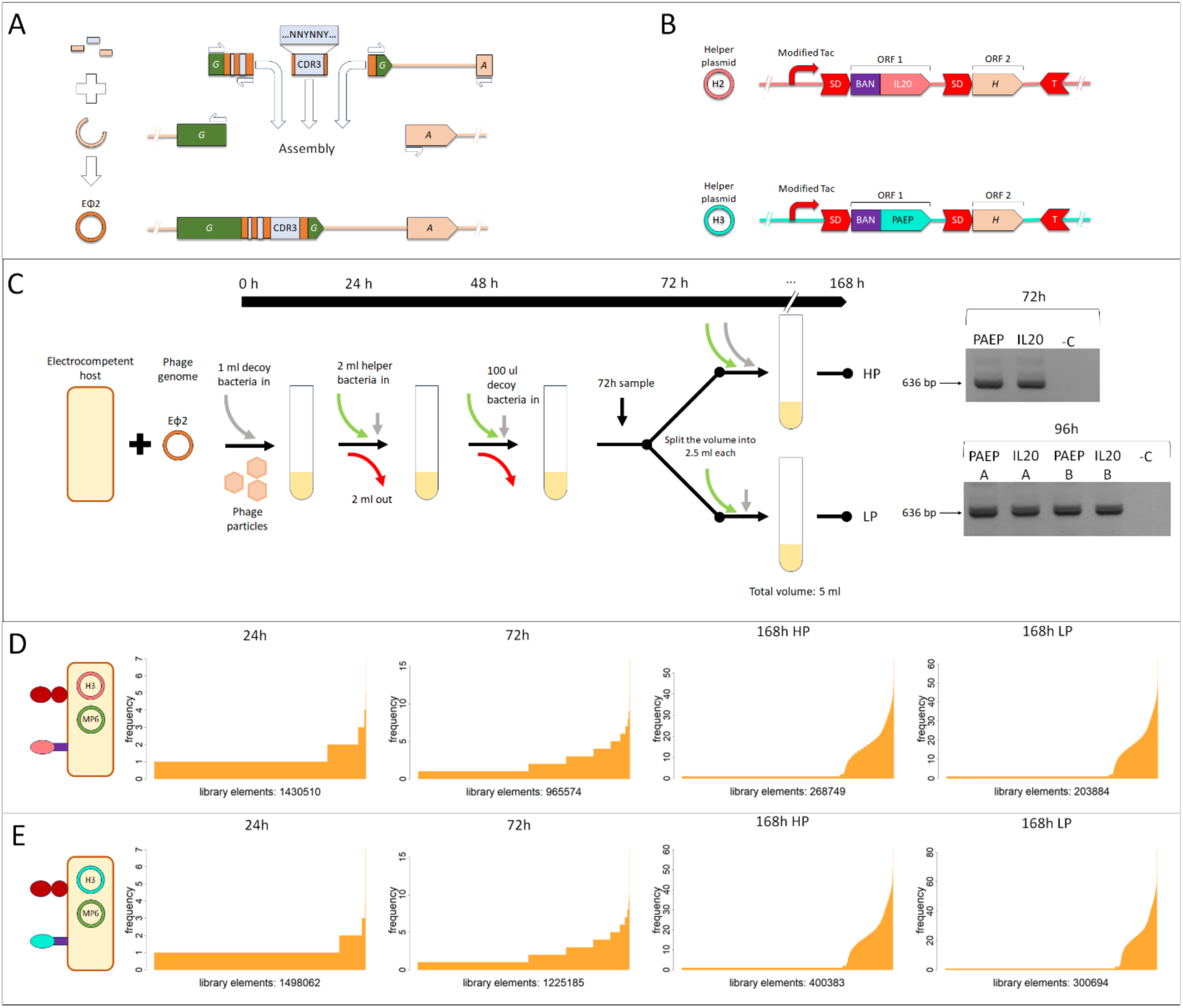
Testing the CDR3 artificial library for two new target proteins. A. Graphical description of the cloning process of engineered phage EΦ2 which correspond to the construction of the CDR3 artificial library. B. Diagram showing the layout of the helper plasmids H2 (IL20RA) and H3 (PAEP). B. The setup of for both experiments is shown, at the 72h time point, the culture was split into 2: high-pressure conditions (HD) and low-pressure conditions (LP). A PCR was performed to check for the presence of the phages after and before the split of the cultures. C. DNA samples at different time points and conditions were taken from the IL20RA cultures and the variant distribution is shown. D. Identical analysis was performed for the PAEP cultures.

EΦ2 was cultured under low selection conditions for H2 and H3 and confirmed by PCR at the previous threshold where propagation was necessary for continued detection, 72h (**Figure 6C**). To better explore optimal conditions for SurPhACE, cultures were split for continued low pressure selection (LP) and high-pressure selection (HP). Both conditions yielded detectable EΦ2 past 96h and finally harvested at 168h.

To expand our range of analysis we conducted long read sequencing, covering the full nanobody and the N-terminal fragment of protein G, but first asked how our random CDR3 library changed during SurPhACE. From 24h post electroporation to the end of the experiment, there was a reduction in library size and enrichment of individual mutants for both targets (**Figure 6C**). CDR3 remained average size of 60 bp in all samples with very variance of (60.8 ± 5.4 bp) (**Supplementary Figure 2**), suggesting a high number of correctly assembled phages. The number of individuals per variant showed a near uniform composition during the first 24h for both targets with enriched variants emerging over time (**Figure 6D and 4E**). Counter to our expectation, LP conditions resulted in a smaller final library size for both targets, however even the most abundant mutant at this time point constituted <0.01% of the library, suggesting longer selection was necessary to arrive at conditions where sequencing was not necessary to detect a dominant mutant.

### Enriched CDR3 variants form subsets of partially converged motifs

To investigate whether library reduction and enrichment was due to selection, or random drift, we examined the 20 most abundant CDR3 variants per treatment, looking for evidence of nonrandom variation and assuming the shared CDR1 and 2 domains would create some target bias (**Figure 7**). Using hierarchical cluster analysis to group similar features, enriched variant subsets showed convergence that could not be explained by chance. The IL20RA library presented a shared motifs of up to 6 identical amino acids that (p=3X10^-16^) (**Figure 7A**), while the PAEP library featured a cluster of 3 shared 3 amino acid motifs (p=8X10^-20^) (**Figure 7B**). In both cases only one variant did not show some sequence similarity to at least one other enriched variant.

**Figure 7.**
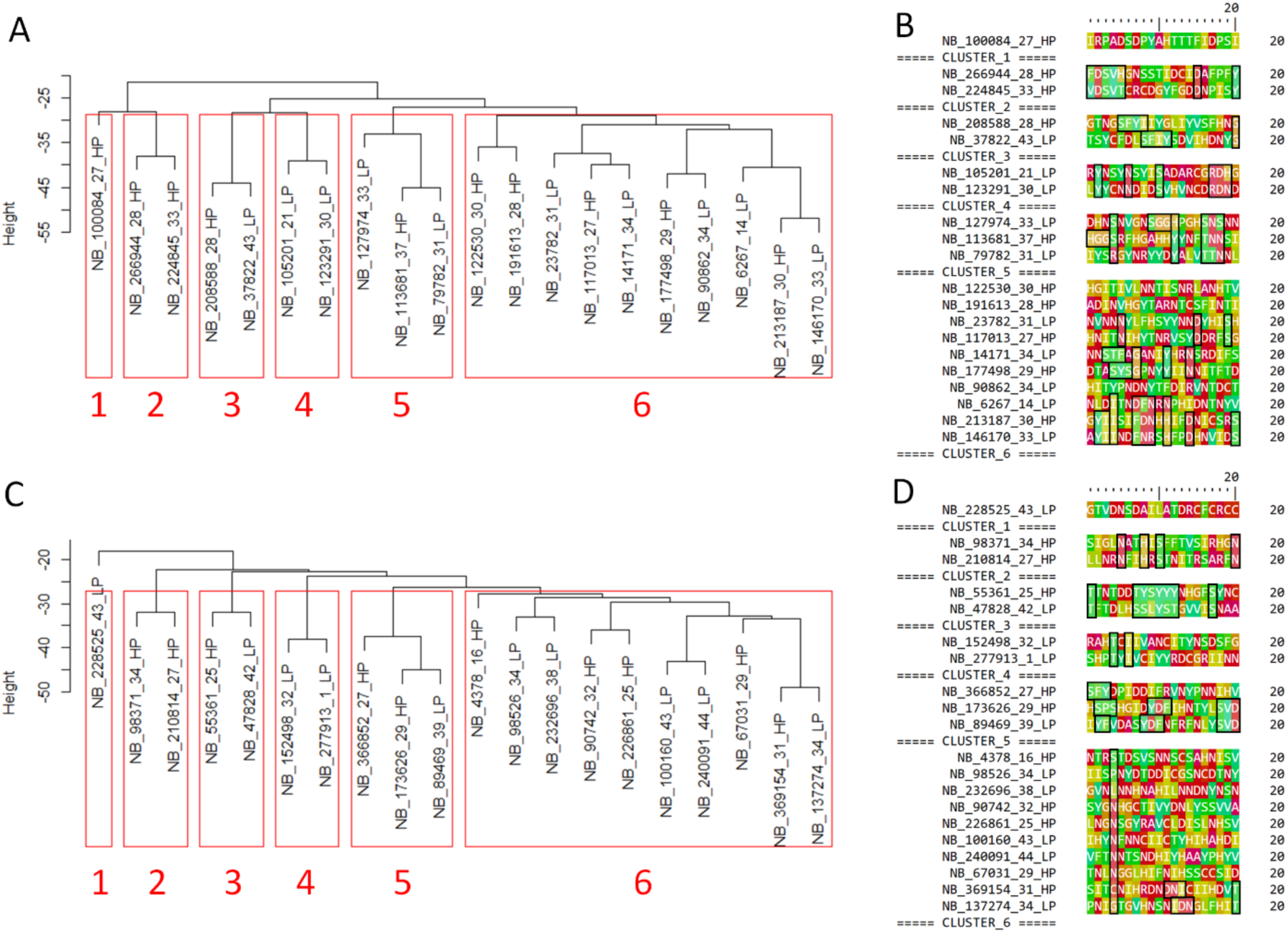
CDR3 Hierarchical Cluster analysis. A. Dendrogram of the 10 most abundant IL20RA variants for each treatment. B. The IL20RA sequences ordered by cluster number, black boxes denote patterns. C. Dendrogram of the 10 most abundant PAEP variants for each treatment. D. The PAEP sequences ordered by cluster number, black boxes denote patterns

### SurPhACE generates large numbers of spontaneous mutations, but truncation mutations were strongly enriched

We took our top 10 enriched variants and traced their lineage in the library back to 24h electroporation to investigate the influence of MP6 induced random mutations across the whole nanobody sequence on library enrichment. Mutations accumulated across the time course as expected, with sub-variants of insertions and deletions (indels) being common feature (**Figure 8A and Figure 8B**). While every enriched CDR3 had a sub-variant that expressed a complete nanobody at the final sampling, multiple indel sub-variants also shared the same CDR3 sequence. In all cases, once an indel sub-variant arose mutations became more common for that sub-lineage compared to full length clones (**Supplementary Figure 4 and Figure 5**). The predominant indel hot spot occurred at the interface between the 5’ nanobody sequence and protein G gene (19-82 bp). Substitution mutations showed minor enrichment at the 5’ end of the sequence and within CDR2 (**Figure 8C**). Overall, the mutational patterns suggested selection pressure for integrating the nanobody linkage or not expressing the nanobody, but not necessarily deleting protein G entirely.

**Figure 8.**
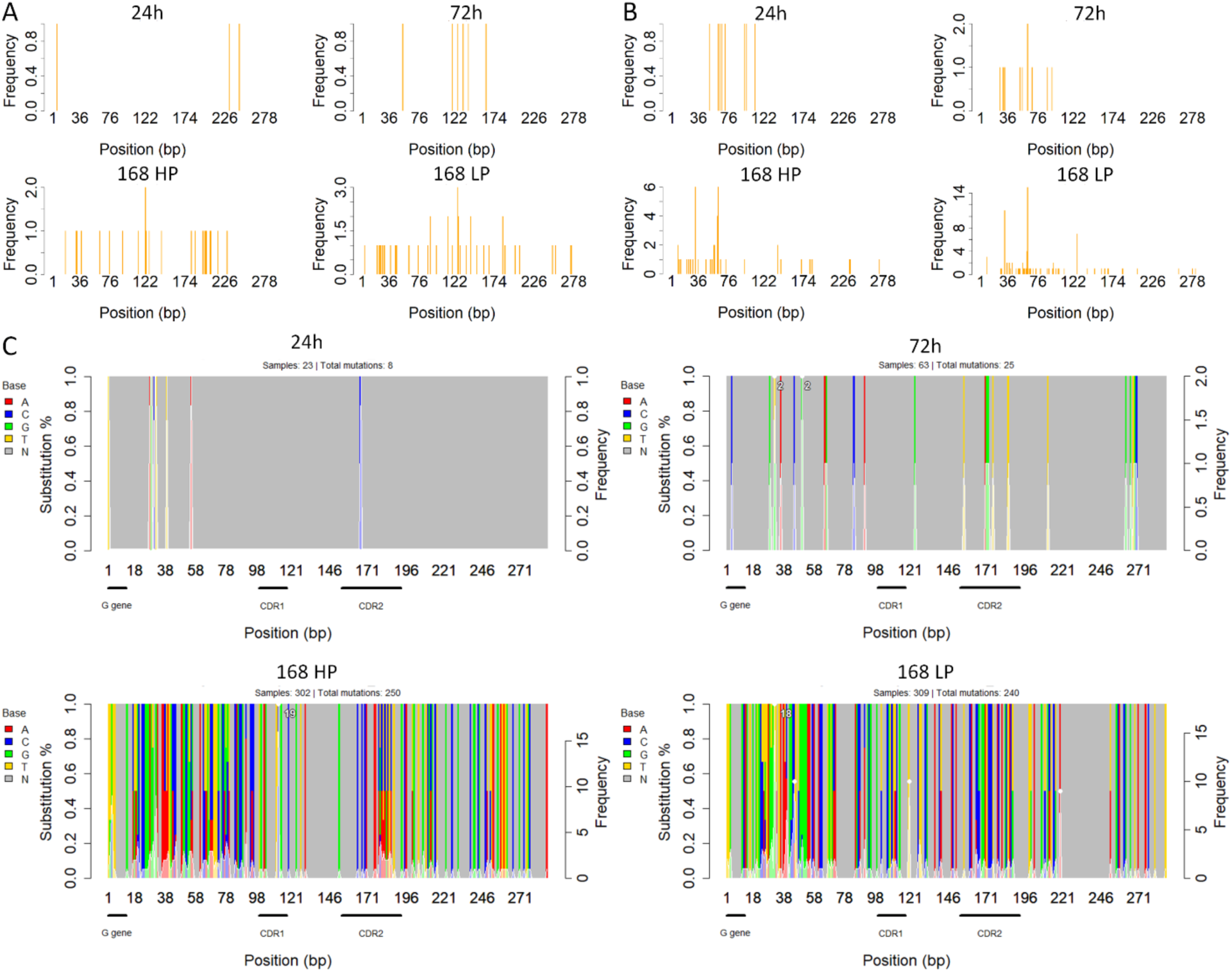
Mutation map of the IL20RA cultures of the 10 most abundant variants excluding CDR3. A. Insertions events at different time points and conditions. B. Deletion events at different time points and conditions. **C**. Substitution map of each time point and condition, the CDR1, CDR2 and part of the G gene are represented. The left Y axis represent the base composition ratio per nucleotide and the right Y axis the mutation density represented by the white line and shaded area. The highest density points are highlighted with a white dot and the maximum peak with the frequency.

### Binding Assays???

#### SurPhACE library assays with FCGR2A protein produced 2 tentatively high affinity binders

The CDR3 sequences of the 10 most abundant candidates from each of the 6 replicates of the FCGR2A libraries (**Supplementary Figure 1**) were put into the nanobody backbone, translated and their interaction with their targets modeled. The properties of these models can be seen on **Table 1**. The candidates binding to FCGR2A had similar properties to the nanobody-αFcR2B - FCGR2B complex, suggesting a strong binding to their target. Surprisingly, these 2 candidates were present in the six replicates, but not necessarily having the higher frequencies of their corresponding replicates

**Table 1.**
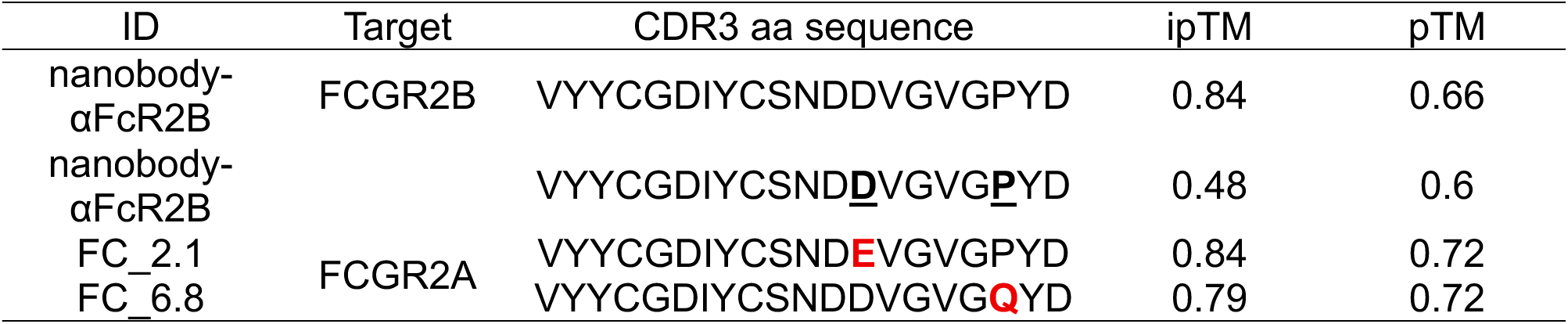
AlphaFold predicted properties of the nanobody-target complex. The original nanobody against FCGR2B is shown for comparison, the intended target and the new target. Finally, the two best mutants where the amino acids in red are the ones different from the original template, these candidates were also present in the other 6 samples.

#### AlphaFold CDR3 Artificial library models have relatively poor binding properties

The first IL20RA 5 nanobody candidates per treatment were translated and evaluated. The ipTM and pTM were below the recommended confidence values for these two parameters. Some candidates like LP_9 and LP_105201 have higher values than the rest but still below the confidence threshold.

## DISCUSION

SurPhACE combines the most useful aspects of DE methods for engineering new binding molecules: phage display, cell surface display, and PACE methods. Making these approaches compatible required innovation for each component. We developed a novel phage for directed evolution, ΦX174, taking advantage of its natural binding directly to the bacterial cell surface. Using the BAN tag, we were able to key selection directly to cell surface binding, while employing decoy bacteria allowed for simultaneous positive and negative selection of phage. Learning from our dynamic systems model led to a further innovation of starting with rudimentary a library of binders, allowing for exploration of a broad initial sequence space with continual refinement through random mutation. With all these elements in play, SurPhACE offers benchtop DE of binding proteins using only 3 plasmids, a PCR generated synthetic phage, and standard molecular biology equipment.

### Dual selective pressure emulates affinity maturation

Directed evolution is based on mimicking natural selection to improve particular evolvable element features. To do this, selective pressures in the system are selected so that the desirable traits are obtained^51^. We implemented 2 selective pressures in our system, one being binding specificity, in the form of the decoy bacteria and helper bacteria. This selective pressure closely emulates the affinity maturation and cell tolerance of mammalian immune systems by encouraging the proliferation and evolution of the best variants and removing the variants that recognize non target structures on the surface of the bacterium (akin to eliminating autoreactive B cells)^52^. The other selective pressure was the dilution ratio of our semicontinuous setup that washed away low affinity binders.

### DE requires nonsurvival for anything but the selection conditions

All the elements of our system were tested and the results matched our predictions. However, the phage still needed the activation of the MP6 plasmid in order to survive for longer (**Figure 3H**). The phage also needed lower dilution ratios to survive suggesting that some element really crucial for replicating effectively had to be fixed. This might suggest that the insertion point of the nanobody in the G protein might not be ideal causing target binding or capsid assembly difficult^26^. The EΦ1 phages disappearing from the cultures so early in the experiments when host bacteria containing H1 plasmids were used (**Figure 3G and Figure 3I)** indicates that the capsid might not has been stable and this had been seen previously with phage display^53,54^. MP6 played a role in the EΦ1 phages survival during the first 2 days of the culture (**Figure 3H**) but was not enough to keep them replicating with helper bacteria H2 and H3 helper bacteria (**Supplementary Figure 3**). So we concluded that MP6 played a role in stabilizing the capsid by mutating the fusion spike protein that was not related to the affinity of the nanobody towards the target.

### Successful DE depends on selective sweeps

Previous PACE methods subject a single clone to random mutation, with the ongoing challenge being the evolutionary distance between the initial gene and the target functionality. We designed our initial experiments accordingly, by initializing with a nanobody that had some potential binding affinity and, based on the necessity of MP6 to phage survival, some adaptation happened, however out limited sequencing could not tell if a selective sweep of some crucial mutation had occurred. Within the sequenced CDR3 region, we saw poor enrichment of mutated variants (**Figure 4**). This phenomena was supported by our models, when the binding affinity of the majority CDR3 was already sufficient for propagation, small gain-of function mutations could not sweep the population under the experimental conditions used (**Figure 4**, **Figure 5C and Figure 5D**). The current fitness and frequency of EΦ1 were too high for selective sweeps to occur within the time frame of the experiment probably due to frequency-dependent interactions, even though strong binders were among the population and across samples (**Table 1**), however further experiments are needed to validate this^55,56^. Conversely, when there was no binding affinity, SurPhACE could not proceed and phage was lost within 2 days.

### SurPhACE continuous selection at high density is prone to clonal interference

To counter frequency-dependent interactions and expand the fitness landscape available to the phages we implemented an artificial library of variants, a well established DE method^30,51^ (**Figure 6A**). The distribution of variants at 24h was nearly uniform as expected from a random synthetic library prior to selective pressure^57^. Reduction of diversity and enrichment of CDR3 variants required time, with only a 1/5 reduction over 168h (**Figure 6D and Figure 6E**). Compare the continuous low selective pressure to methods like biopanning, where the majority of a library can be washed away in a single step^25^. We suspect this difference is due to the potential of multiple infection events to partially decouple genotype from phenotype. This clonal interference, is a phenomenon observed in large populations of asexual species, that can slow the rate of a selective sweep from progressing through the culture^58–60^.

### High genetic drift during SurPhACE can lead to loss-of-function sub-variants

Genetic drift allows for effective exploration of mutational space and is a precondition to the appearance of many beneficial mutations across the phage, but random effects can also drive the enrichment of negative traits^61,62^. The 10 most abundant variants for each treatment, independent of target, all had multiple loss-of-function indel sub-variants. Given the high culture density these could persist if consistently packaged in hybrid viral particles during multiple infection events (**Supplementary Figure 4 and 5**)^63,64^. These persistent satellite viruses also explain the low enrichment of variants observed, since parasitism would negatively affect the host population and reduce the number of gain-of-function sub-variants (**Table 2**)^65,66^.

**Table 2.**
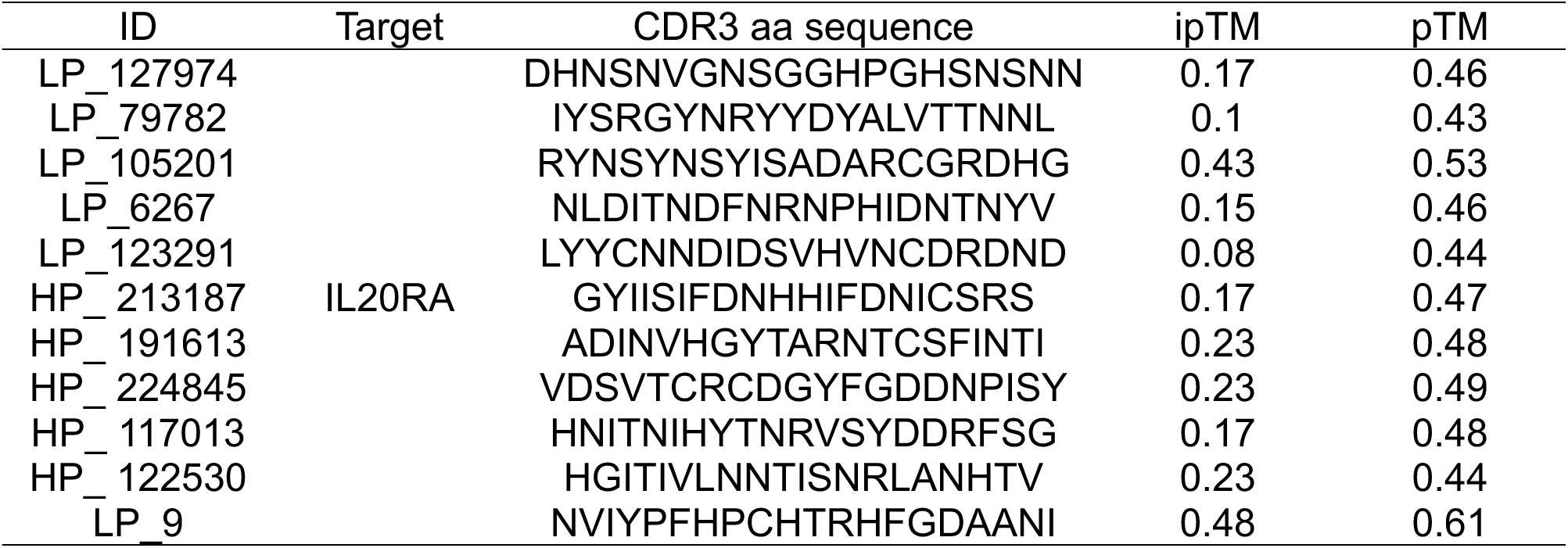
AlphaFold predicted properties of the nanobody-target complex for the CDR3 artificial library. The properties of the 5 more abundant variants per condition (low and high pressure) are shown. LP_9 is a sequence obtained directly from a PCR of the low-pressure library.

When tracing variant lineages, we found that the number of subsequent mutations was greater in the indel sub-variants than full length relatives (**Figure 8**), indicating relaxed selection on downstream sequences and reduced efficiency of positive selection^67–69^. In these indel sequences, genetic drift is the dominant factor for their frequency in the culture. Regardless of target, or when indels occurred across the experiment the linker region between protein G and the nanobody was a mutational hot spot, suggesting some specific role of that region for the phage fitness or a suboptimal positioning of the nanobody in the G protein sequence^57^.

### CDR3 convergence could be due to the common CDR1-2

Among the 20 most abundant variants for each target, some shared potential motifs, demonstrating convergence (**Figure 7**). While it would be statistically improbable that random sequences would optimize for the same epitopes, it is important to note that the whole library shared a common CDR1-2, and was subject to whatever initial affinity these regions had. This is actually a feature of naturally derived antibody libraries as well in that CD1-2 mature prior to the final antigen presentation and education of B-cells. In the spleen, this may be a mechanism to reduce autoimmune affinity, but in our case it simply influences initial binding epitopes. In the future this is a feature that could be leveraged in that a human derived library of CD1-2 variants could have an improved safety profile compared to a truly random library.

SurPhACE presents us with a platform to take advantage of decades of innovations in library design and benchtop DE methods. Future iterations of the method will tune the many details that could optimize SurPhACE from a novel technique to a robust source of high affinity binding molecules. These include dynamic selection pressure to keep variants at the threshold of survival. Tapered mutational rates would allow improved negative selection of indel sub-variants. Normalization steps would enrich rare variants and allow for faster selective sweeps of high binders. Finally, an optimized starting library for given applications would narrow the search space and potentially accelerate initialization to high binder sweeps.

## Supplementary material

**Table 1:** Primers used.

**Table 2.**
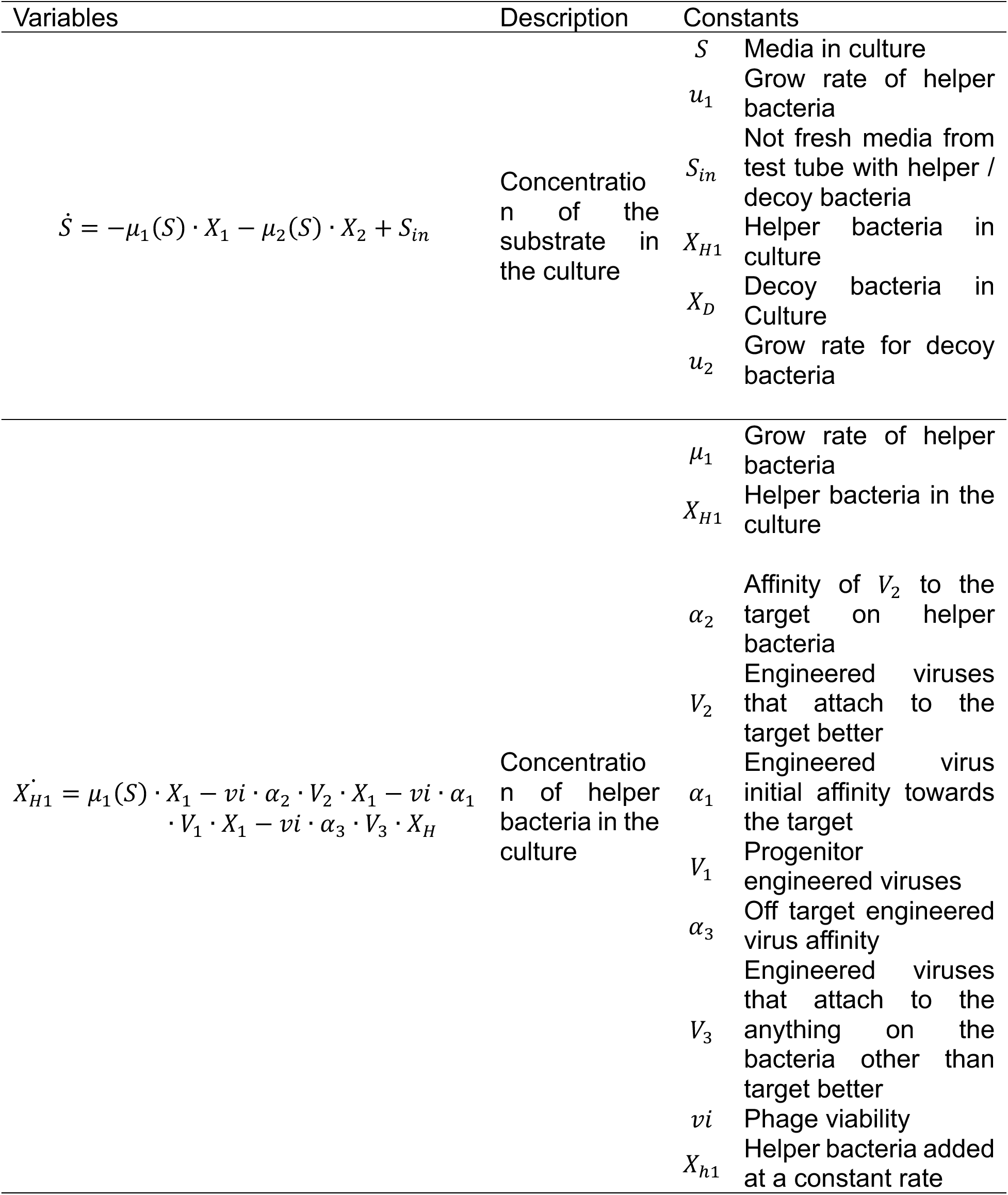
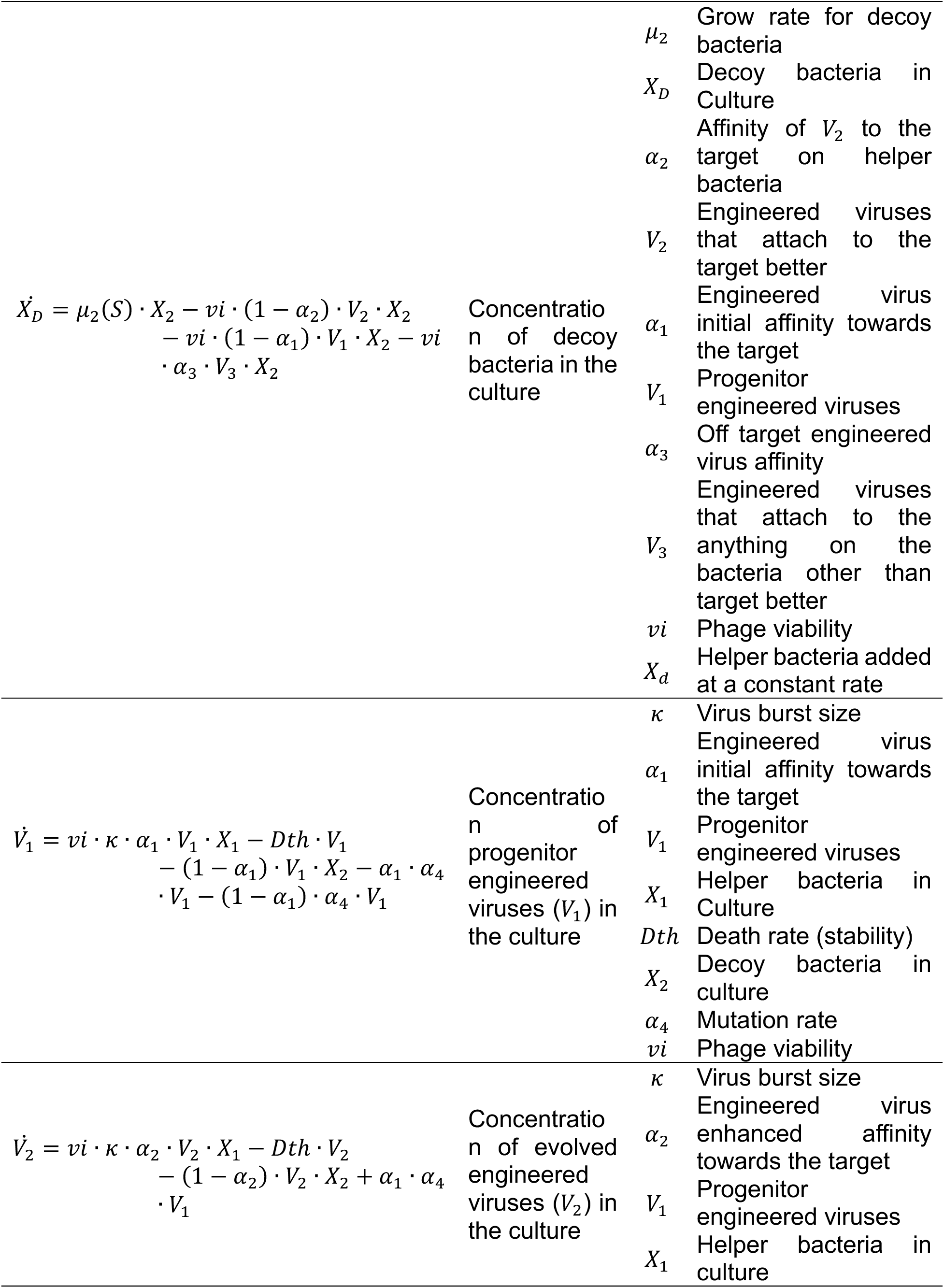
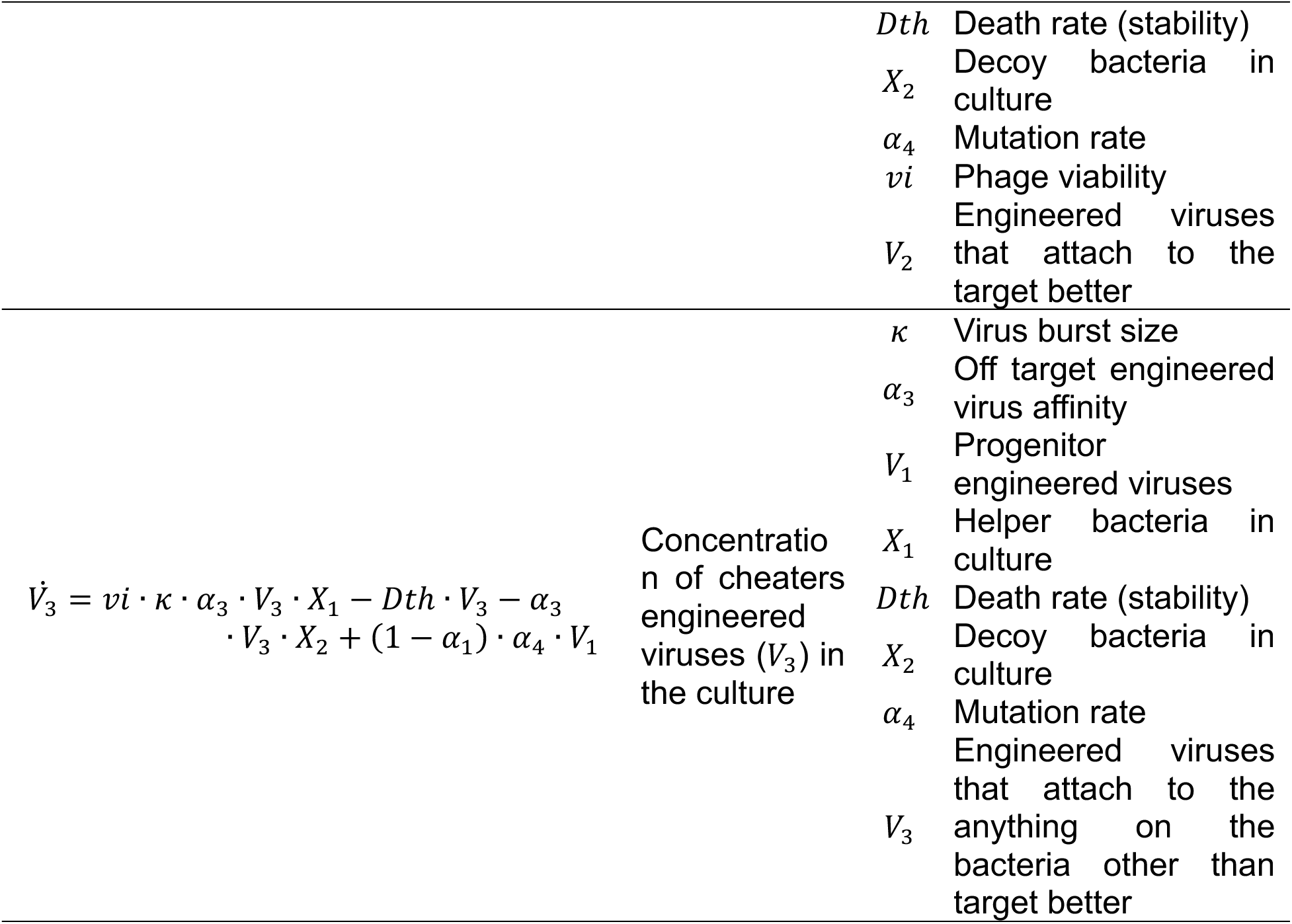
Variables and constants description, units still to be defined. Variables Description Constants.

**Supplementary Fig1.**
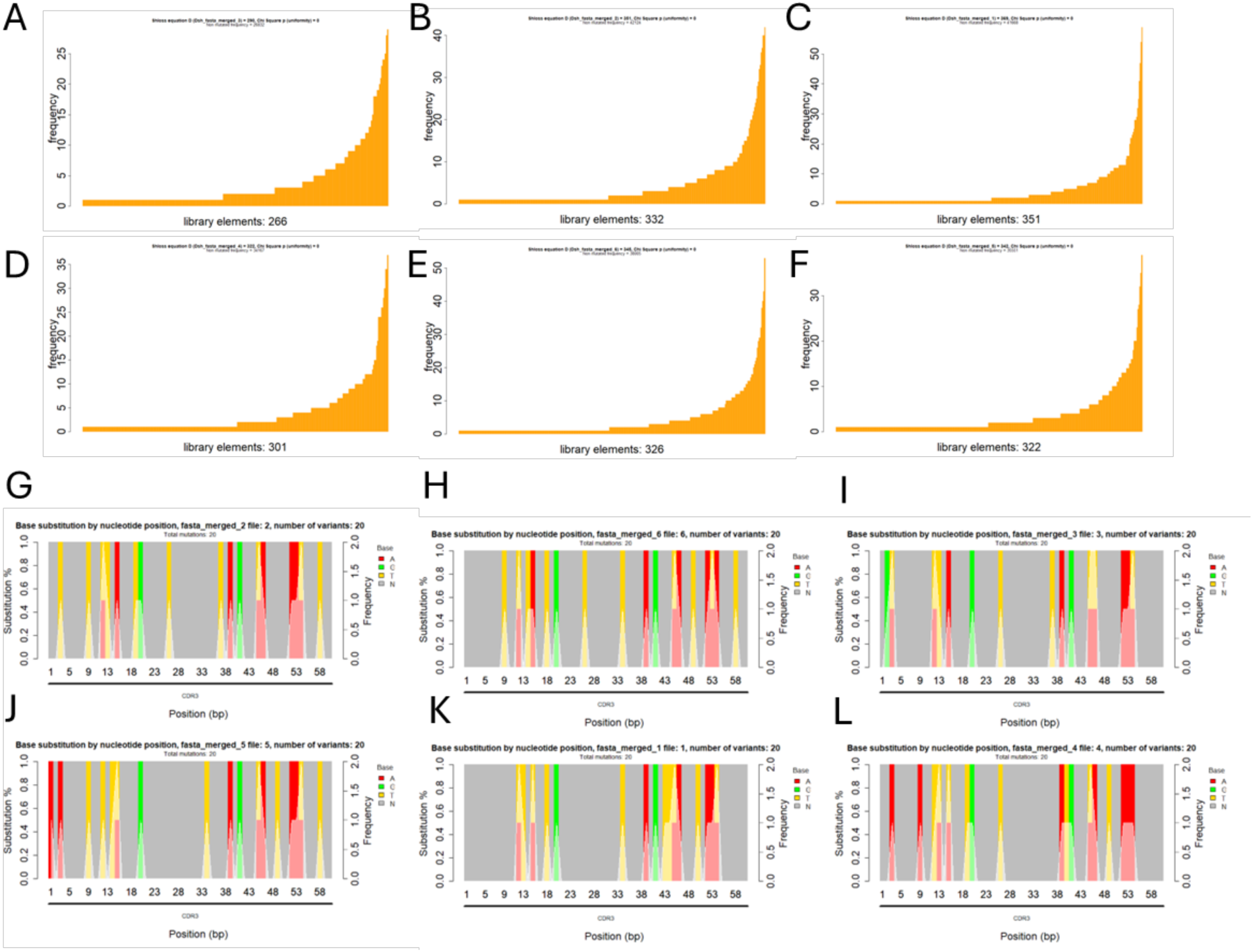
NGS analysis of the 6 independent samples. (**EΦ1 CDR3)**. **A-F.** Number and frequency of CDR3 variants. **G-L** Bar plot showing the substitution percentage (colored bars, N stands for unaltered nucleotide) and mutation density (white line) for the 20 most abundant mutated variants.

**Supplementary fig 2.**
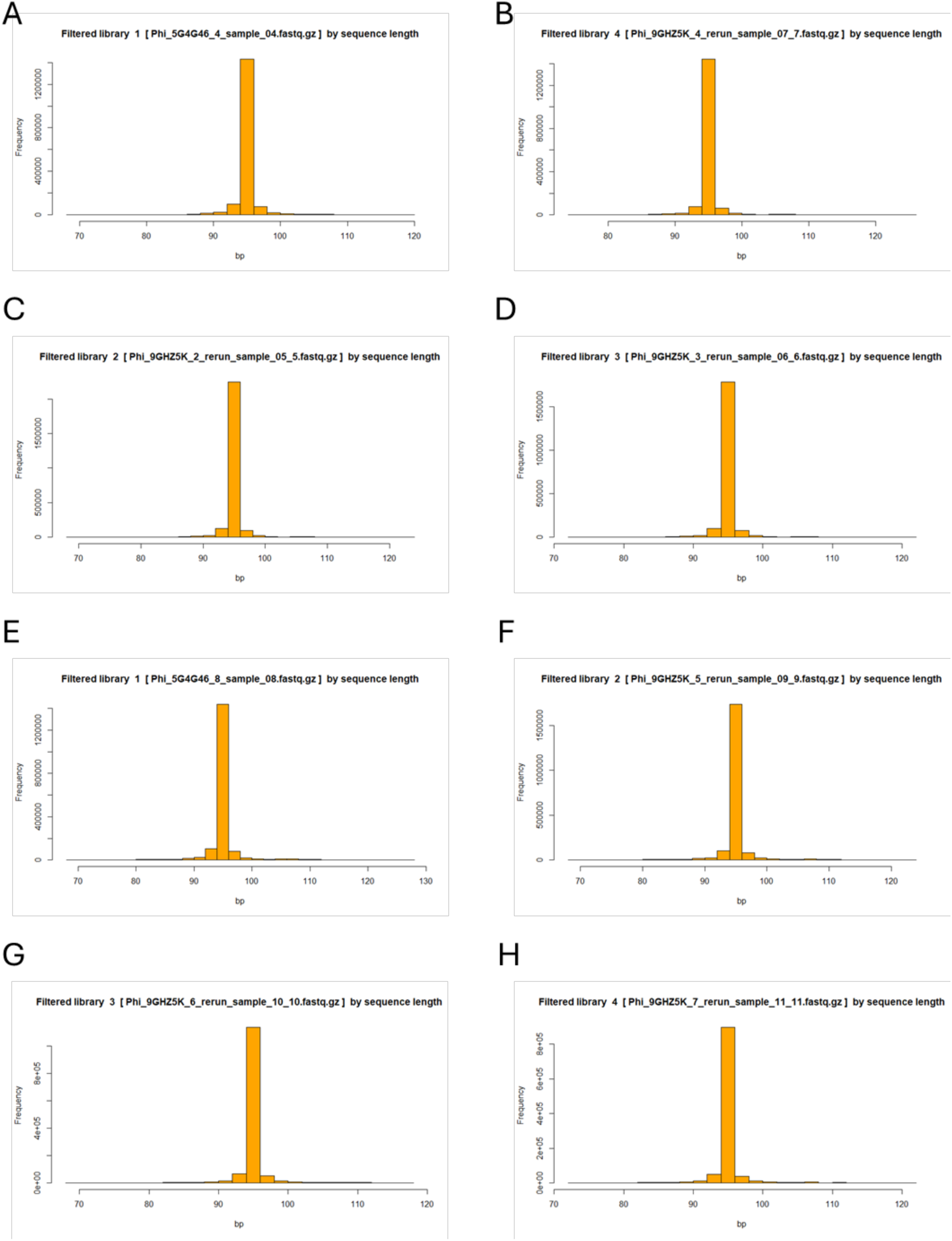
CDR3 size histogram of the libraries (96 bp, 60 bp without flanking sequences^1^. **A-D.** PAEP **E-H.** IL20. All CD3 showed a consistent length during the experiment independently from the target protein. ^1^Flanking sequences were used to map all the CDR3s because they were well conserved among all variants

**S fig 3.**
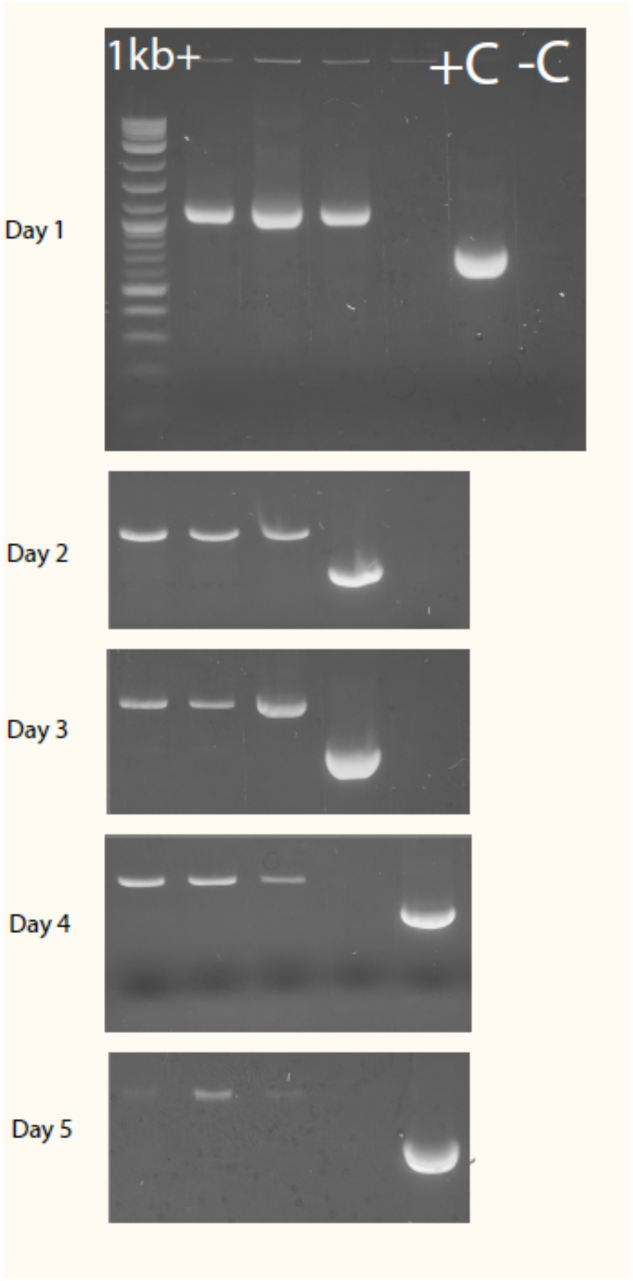
Three replicates showing the EΦ2 fading out of the H2 helper bacteria (IL20RA / MP6) cultures over the course of several days. No decoy was added

**S Fig 4.**
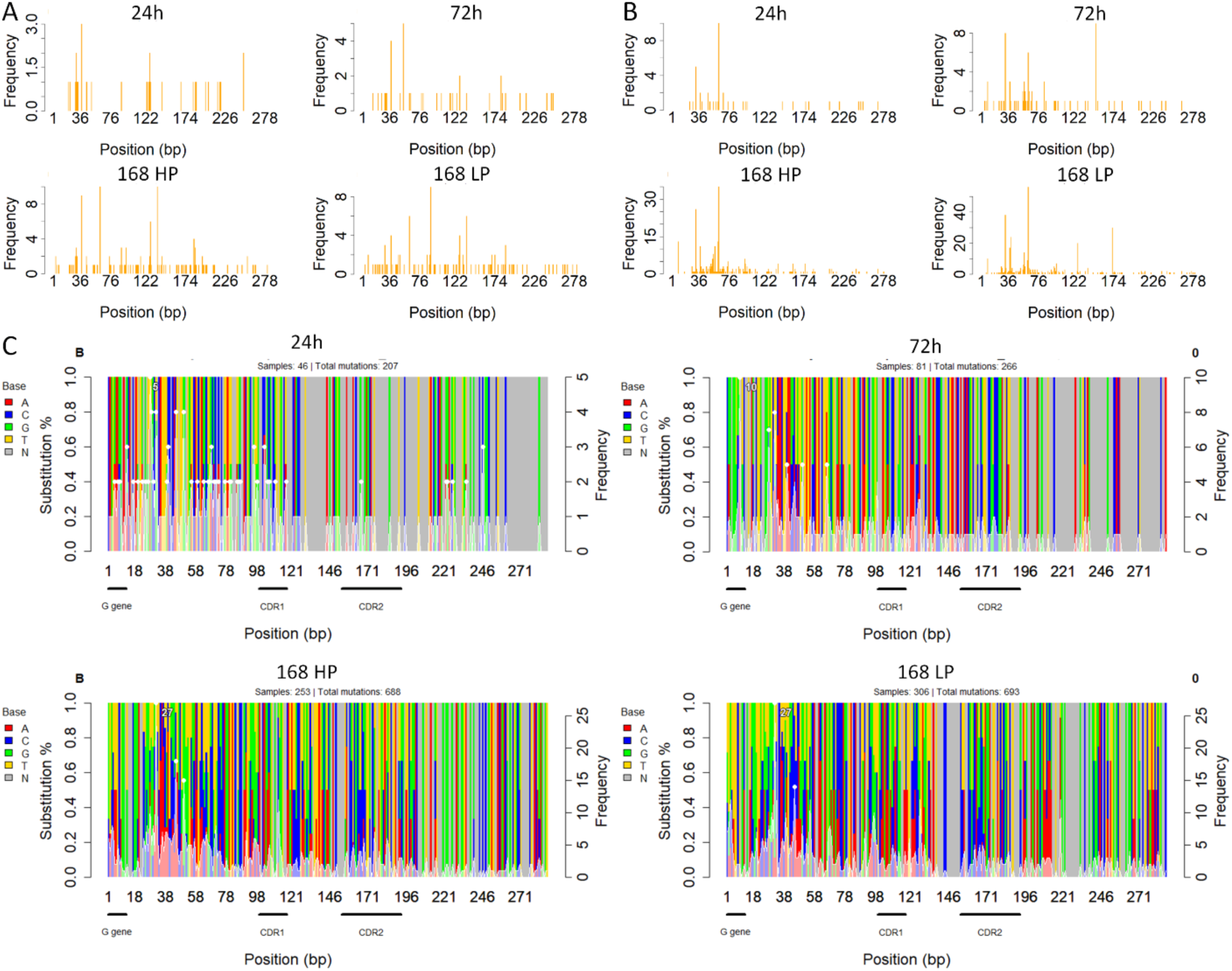
IL20RA.

**S Fig 5.**
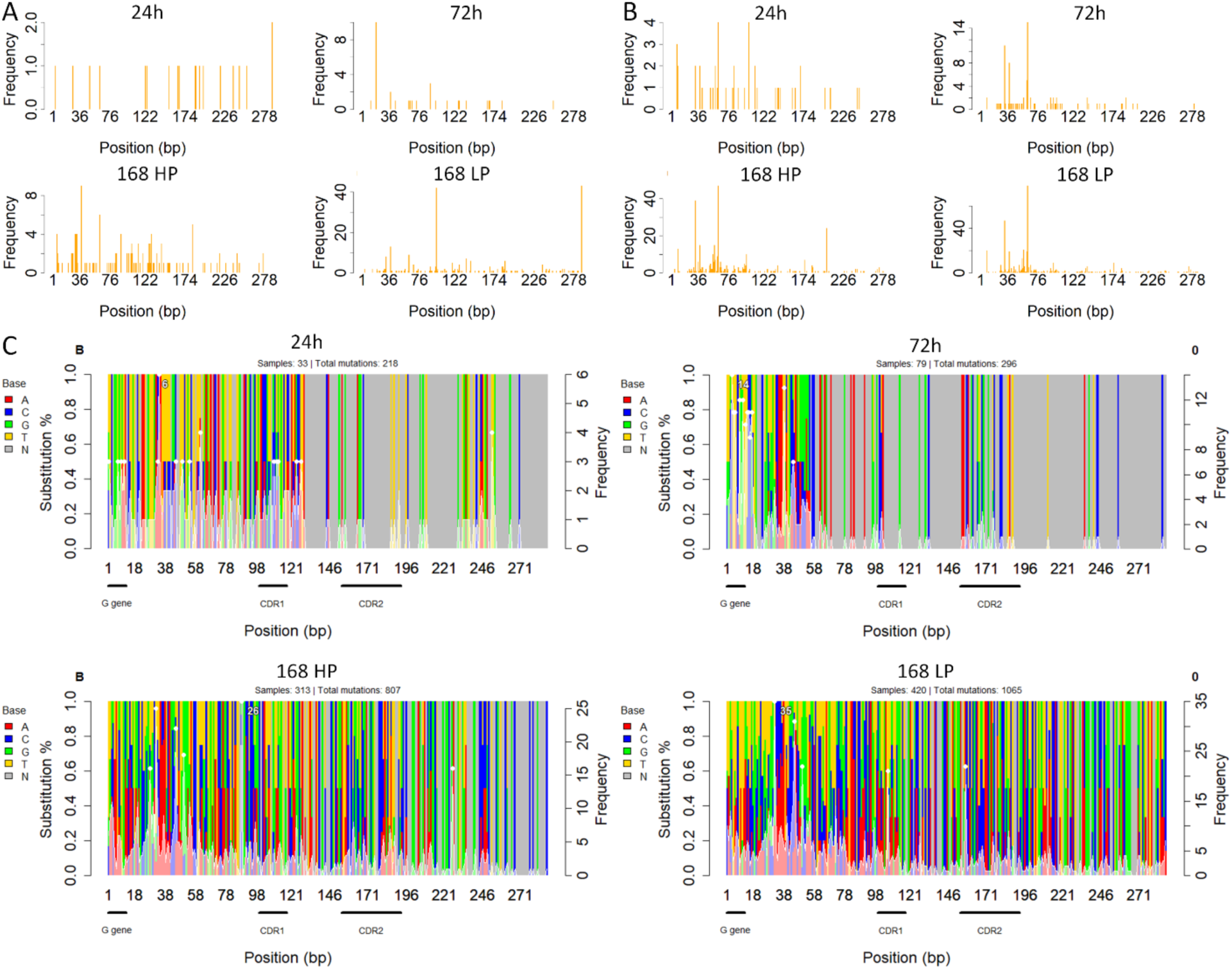
PAEP.

